# Identifying Patients with Cognitive Motor Dissociation Using Resting-state Temporal Stability

**DOI:** 10.1101/2022.10.17.512475

**Authors:** Hang Wu, Qiuyou Xie, Jiahui Pan, Qimei Liang, Yue Lan, Yequn Guo, Junrong Han, Musi Xie, Yueyao Liu, Liubei Jiang, Xuehai Wu, Yuanqing Li, Pengmin Qin

## Abstract

**Background:** Using task-dependent neuroimaging techniques, recent studies discovered a fraction of patients with disorders of consciousness (DOC) who had no command-following behaviors but showed a clear sign of awareness, which was defined as cognitive motor dissociation (CMD). Although many efforts were made to identify the CMD, existing task-dependent approaches might fail when patients had multiple cognitive function (e.g., attention, memory) impairments, and thus lead to false-negative findings. However, recent advances in resting-state fMRI (rs-fMRI) analysis allow investigation of the dynamic change of spontaneous brain activity, which might be a powerful tool to test the patient’s cognitive functions, while its capacity in identifying CMD was unclear.

**Methods:** The rs-fMRI study included 119 participants from three independent research sites. A sliding-window approach was used to investigate the dynamic functional connectivity of the brain in two aspects: the global and regional temporal stability, which measures how stable the brain functional architecture is across time. The temporal stability was compared in the first dataset (36/16 DOC/controls), and then a Support Vector Machine (SVM) classifier was built to discriminate DOC patients from controls. Furthermore, the generalizability of the SVM classifier was tested in the second independent dataset (35/21 DOC/controls). Finally, the SVM classifier was applied to the third independent dataset where patients underwent an rs-fMRI and brain-computer interface assessment (4/7 CMD/potential non-CMD), to test its performance in identifying CMD.

**Results:** Our results showed that the global and regional temporal stability were impaired in DOC patients, especially in regions from the cingulo-opercular task control, default mode, fronto-parietal task control, and salience network. Using the temporal stability as features, the SVM model not only showed a good performance in the first dataset (accuracy = 90 %), but a good generalizability in the second dataset (accuracy = 82 %). Most importantly, the SVM model generalized well in identifying CMD in the third dataset (accuracy = 91 %).

**Conclusion:** The current findings suggested that rs-fMRI could be a potential tool to assist in diagnosing CMD. Furthermore, the temporal stability investigated in this study also contributed to a deeper understanding of the neural mechanism of the consciousness.

## Introduction

Consciousness is a fundamental aspect of our complex mental life, of which the underlying neural mechanism has been a topic of great interest in the area of neuroscience.^1,2^ In the clinical setting, severe brain injury would lead to the disorders of consciousness (DOC), where some patients showed no awareness of the environment or of themselves (i.e., vegetative state, VS, also called unresponsive wakefulness syndrome, UWS),^3^ some showed unstable but reproducible signs of awareness (i.e., minimally conscious state, MCS)^4^ that could be assessed by their behavioral responses.^5^ To date, the diagnosis of patients with DOC was largely dependent on their behavioral response at the bedside, while this indirect approach may bias the evaluation of the patients’ actual awareness due to their cognitive or sensory impairments.^6,7^ Thus, many efforts were made to detect patients’ actual awareness by means of neuroimaging (EEG and fMRI) and brain-computer interface (BCI) techniques with specific active tasks, where a fraction of (around 20%) DOC patients were found able to follow commands and show a similar brain activation pattern as healthy controls.^8–11^ These patients were defined as cognitive motor dissociation (CMD), also known as the functional locked-in syndrome.^12^ However, current tools for identifying CMD requires that patients could understand and follow the researchers’ oral command in order to perform a specific task, which may fail due to the patients’ cognitive function (e.g., attention, memory) impairments, thus leading to a false-negative finding.^13^ To this end, as a convenient (i.e., typically acquiring enough data in less than 10 minutes) and task-free (i.e., no active participation required) neuroimaging technique, resting-state fMRI (rs-fMRI) accompanied with the BCI technique, could be an ideal complementary approach to identify CMD. However, evidence for its diagnostic value is still scarce.

Using static rs-fMRI functional connectivity (FC), plenty of studies have observed that a number of functional networks were impaired or reorganized in DOC, which were associated with decreased awareness. These networks include the default mode network,^14,15^ sensorimotor network,^16,17^ frontal parietal network,^18,19^ etc. However, increasing evidence showed that the functional networks in the brain were not stationary, and researchers therefore combined different approaches, such as dynamic functional connectivity (dFC) and the clustering method, and identified a number of discrete and reproducible brain states.^20^ For instance, studies have shown that altered brain dynamics (e.g., dwelling time, pattern transition) were associated with the DOC, which was not found with conventional FC approaches based on a stationarity assumption.^21–23^ Although, most of the existing findings on the DOC investigated the dFC properties from the perspective of the functional organization of the whole-brain, while the dynamic FC pattern of a given region hasn’t been sufficiently studied. For this, temporal variability, an alternative dFC measurement approach was recently developed. This approach does not cluster brain activity into several discrete brain states, but explores the global and regional functional dynamics in a more straightforward way.^24,25^ According to its definition, a brain region with high temporal variability indicates lower temporal stability (i.e. regional activity being less stable across time), and vice versa.^24,25^ Using these temporal properties, studies have found that alteration of the temporal stability of certain brain regions could be highly informative for identifying various mental disorders with cognitive function impairments, e.g., Alzheimer’s disease, Schizophrenia, and autism.^25–27^ However, it is unclear whether and how the temporal stability altered in patients with DOC, and more importantly, whether this measure could be used to correctly identify CMD.

The aim of this study was twofold: first, to investigate the difference of temporal stability between DOC and controls and build a linear support vector machine (SVM) classifier, and second, to test the performance of the SVM model in correctly identifying the patients with CMD. For that, a final cohort of 119 participants were recruited from three independent centers, and divided into three datasets. For our first aim, the first dataset (n = 52, 36 DOC) was used to investigate the difference of temporal stability between the DOC and controls, for which an SVM classification model was built to discriminate DOC patients from controls. The SVM model was then applied to the second dataset (n = 56, 35 DOC) to test its generalizability. For our second aim, the third dataset (n = 11, 4 was subsequently diagnosed as CMD) was used, to further test the SVM model’s generalizability. Our results first showed that the global and regional temporal stability were indeed impaired in the patients with DOC compared with the controls. More importantly, the SVM model not only classified DOC and controls in the first and second datasets with a good performance (accuracy = 90 %, 82 %, respectively), but showed a good generalizability for identifying patients with CMD (accuracy = 91 %) in the third dataset.

## Materials and methods

### Participants

A cohort of 128 participants were recruited from three independent research sites: Huashan Hospital in Shanghai, Zhujiang Hospital in Guangzhou, and Guangzhou General Hospital of Guangzhou Military Command. The above cohort was divided into three datasets: (1) dataset DOC-HS (n = 58, all participants were recruited from Huashan Hospital, note that some of patients were previously analyzed and published out of a different purpose^16,28^); (2) dataset DOC-ZJ (n = 59, all participants were recruited from Zhujiang Hospital); (3) dataset CMD-GH/ZJ (n = 11, 8 participants from Guangzhou General Hospital of Guangzhou Military Command, 3 participants from Zhujiang Hospital). The first two datasets (DOC-HS, DOC-ZJ) were collected for the first aim of the current study. Specifically, dataset DOC-HS was analyzed and reported as the main results, which was also used as a training dataset for the classification analysis, and dataset DOC-ZJ served as an independent dataset for testing. The third dataset (CMD-GH/ZJ) was collected for the second aim of the study. For these patients, inclusion criteria were: (1) diagnosed as having UWS or MCS using Coma Recovery Scale–Revised scale;^5^ (2) scanned at least 4 days after the acute brain insult; (3) had structurally well-preserved brain images that were carefully chosen by author XW and checked by author PQ. Patients were excluded due to contraindication for MRI, and motion artifacts (i.e., > 20% volumes were identified as outliers in preprocessing) during MRI scanning.

According to the exclusion criteria, a total of 9 participants (6 from DOC-HS, 3 from DOC-ZJ) were excluded due to motion artifacts, resulting in the final cohort of 119 participants (please see Table 1 and Supplementary Tables 1 - 3 for detailed demographic and clinical characteristics of each dataset). Specifically, dataset DOC-HS included 36 DOC patients (UWS/MCS: 26/10, 29 males, mean age 43 ± 14 years, 25 traumatic, 11 non-traumatic, mean time since brain injury: 85 days) and 16 fully conscious participants with a history of brain injuries (10 males, mean age 40 ± 17 years, 12 traumatic, 4 non-traumatic, mean time since brain injury: 97 days). Dataset DOC-ZJ included 35 DOC patients (UWS/MCS: 26/9, 21 males, mean age 43 ± 15 years, 13 traumatic, 22 non-traumatic, mean time since brain injury: 74 days) and 21 healthy controls (10 males, mean age 32 ± 10 years). Dataset CMD-GH/ZJ included 11 DOC patients (UWS/MCS: 8/3, 4 CMD; 9 males; mean age 45 ± 13 years, 5 traumatic, 6 non-traumatic, mean time since brain injury: 78 days). All healthy participants recruited in the current study reported no history of neurological or psychiatric disorders. Informed written consent was obtained from all healthy participants, and legal representatives of the patients. The study was approved by the Ethics Committee of the Huashan Hospital, the Zhujiang Hospital, and the Guangzhou General Hospital of Guangzhou Military Command, respectively.

**Table 1.** Demographic and Clinical Information for the Three DOC Datasets

| Dataset | Group | Sample Size | Mean Age (SD) | Sex | Days Since Injury (SD) | Traumatic Injury | Diagnosis (UWS/MCS) | CMD |
| --- | --- | --- | --- | --- | --- | --- | --- | --- |
| DOC-HS | BI | 16 | 40 (17) | 10M/6F | 97 (153) | 12 | - | - |
|  | DOC | 36 | 43 (14) | 29M/7F | 85 (87) | 25 | 26/10 | - |
| DOC-ZJ | HC | 21 | 32 (10) | 10M/11F | - | - | - | - |
|  | DOC | 35 | 43 (15) | 21M/14F | 74 (56) | 13 | 26/9 | - |
| CMD-GH/ZJ | CMD | 11 | 45 (13) | 9M/2F | 78 (60) | 5 | 8/3 | 4 |

Note: For the dataset DOC-HS and DOC-ZJ, BI and HC were both considered as controls, whose awareness were intact compared with the DOC patients. For the dataset CMD-GH/ZJ, 11 DOC patients underwent both the resting-state fMRI scanning and brain-computer interface experiment. BI = fully conscious participants with a history of brain injury; DOC = disorders of consciousness; HC = healthy controls; CMD = cognitive motor dissociation; UWS = unresponsive wakefulness syndrome; MCS = minimally conscious state; SD = standard deviation. “-” means not applicable.

### Brain-Computer Interface Procedure

For the 11 DOC patients in the dataset CMD-GH/ZJ, an EEG-based BCI experiment was carried out after the rs-fMRI scanning to identify patients with CMD. These patients were analyzed and reported in a previous study out of a different research purpose.^11^ In the BCI experiment, the participants were instructed to focus on the target stimulus and perform simple tasks. Specifically, a calibration session and an online evaluation session was presented for each patient, where the first session contained 10 trials for training an SVM classifier to detect photograph-related P300, and the second session contained five blocks of 10 trials each, to update the SVM model obtained from the first session. To detect residual awareness in the DOC patients, three different stimuli were adopted, including audiovisual, numbers, and photograph stimuli. The reason we used different stimuli was because of the heterogeneity of the symptoms, where some patients may retain intact visual and auditory function (i.e., suitable for audiovisual stimuli), some may only retain visual function (i.e., suitable for photograph and number stimuli). For the photograph paradigm, the patients were asked to focus on photographs of their own face or strangers’ face in a random order. Before each trial, an audiovisual instruction in Chinese was presented: ‘Focus on your own photograph (or the stranger’s photograph) and count the flashes of the photo frame’, which lasted for 8 s and indicated the target photograph. Then, two photographs were displayed, one of them had a flashing flame, which was randomly selected and flashed five times. After 10 s, one of the two photographs identified by the BCI algorithm was displayed in the center of the screen as feedback, where patients were informed whether they selected the target photograph correctly. The experimental procedure for the number or audiovisual paradigm was similar to the photograph paradigm except that the photographs were replaced by numbers or audiovisual stimuli, respectively.

Each patient was assigned one of the three paradigms, as selected by their family members. Among the 11 patients in the current study, 8 patients participated in the photograph paradigm, 2 in the number paradigm, and 1 in the audiovisual paradigm. Finally, a DOC patient would be diagnosed with CMD when his/her BCI accuracy was higher than 64%, determined at a significance level of p = 0.05 using a χ^2^ test. For detailed information of the BCI experiment, please see the reference paper.^11^

### MRI acquisition

For participants recruited from the Huashan Hospital, the MR images were acquired using a Siemens 3 Tesla scanner. Functional images were acquired using a T2*-weighted EPI sequence (TR/TE/θ = 2000ms/35ms/90°, FOV = 256 × 256 mm, matrix = 64 × 64, 33 slices with 4-mm thickness, gap = 0 mm, 200 scans). For participants recruited from the Zhujiang Hospital, the MR images were acquired using a Philips Ingenia 3 Tesla scanner. Functional images were acquired using a T2*-weighted EPI sequence (TR/TE/θ = 2000ms/30ms/90°, FOV = 224 × 224 mm, matrix = 64 × 64, 33 slices with 3.5-mm thickness, gap = 0.7 mm, 240 scans). For participants recruited from the Guangzhou General Hospital of Guangzhou Military Command, the MR images were acquired using a GE signa 3 Tesla scanner. Functional images were acquired using a T2*-weighted EPI sequence (TR/TE/θ = 2000ms/30ms/90°, FOV = 240 × 240 mm, matrix = 64 × 64, 35 slices with 4-mm thickness, gap = 0 mm, 240 scans). For each participant, a high-resolution T1-weighted anatomical image was also acquired for functional image registration and localization.

### Preprocessing

For all participants across the three datasets, the same preprocessing analysis procedures were performed, using the CONN toolbox (CONN; http://www.nitrc.org/projects/conn, version 21.a) based on the Statistical Parametric Mapping (SPM) 12 program (http://www.fil.ion.ucl.ac.uk/spm, version 7771) implemented in MATLAB 2019a. For each participant, a standard preprocessing pipeline was adopted, following recent studies^29,30^, which included the following steps: (1) removal of the initial four functional volumes; (2) functional realignment and motion correction; (3) functional slice-timing correction; (4) identification of outlier volumes for subsequent regression using the artifact detection tool (ART, http://www.nitrc.org/projects/artifact_detect), in which the default CONN settings of 5 global signal Z-values and 0.9 mm was chosen; (5) spatial normalization into standard Montreal Neurological Institute (MNI) space (resampled to 3 mm isotropic spatial resolution); (6) segmentation of functional and structural data into grey matter, white matter, and CSF tissue; (7) spatial smoothing using a Gaussian kernel of 6mm full-width at half-maximum. In each step, the processed functional and anatomical images were carefully checked with visual inspections. After the preprocessing steps, 9 participants with over 20% outlier volumes identified with ART were excluded from the subsequent analysis (6 from dataset DOC-HS, 3 from dataset DOC-ZJ).

To further reduce cardiac and motion artifacts, the anatomical component-based noise correction method (aCompCor)^31^ was applied to denoise the functional data, as implemented in CONN toolbox was applied to denoise the functional data. Specifically, the aCompCor method denoises functional data by regressing out several potential confounding effects: five potential noise components from the white matter and cerebrospinal fluid signals, estimated motion parameters (3 translation and 3 rotation parameters plus their associated first-order derivatives), the artifacts identified by ART; the main effect of scanning condition. Finally, a linear detrending was applied, following a temporal band-pass filtering between 0.01 and 0.1 Hz, to minimize both low-frequency drift effects and high-frequency noise. The above denoising procedures have been used by several recent rs-fMRI studies that focused on DOC, which were proved to be advantageous in keeping the temporal integrity of the data.^32^ Please see Supplementary Figure 1 for the effect of the denoising procedures on an example.

### Definition on Regions of Interest

The pipeline of data analysis was shown in Figure 1. A well-established network template was used to be slightly modified according to previous studies^33^, to obtain the regional time series. As shown in Figure 1A, this template contained 226 functional regions (5 mm radius spheres, 33 voxels per sphere), each assigned to one of 10 functional networks: auditory (AN), cingulo-opercular task control (COTC), default mode (DMN), dorsal attention (DAN), fronto-parietal task control (FPTC), salience (SAN), sensory/somatomotor (SMN), subcortical (Sub), ventral attention (VAN), and visual networks (VN). For each region of interest (ROI), time series was obtained by averaging the signals of every voxel within the ROI.

**Figure 1.**
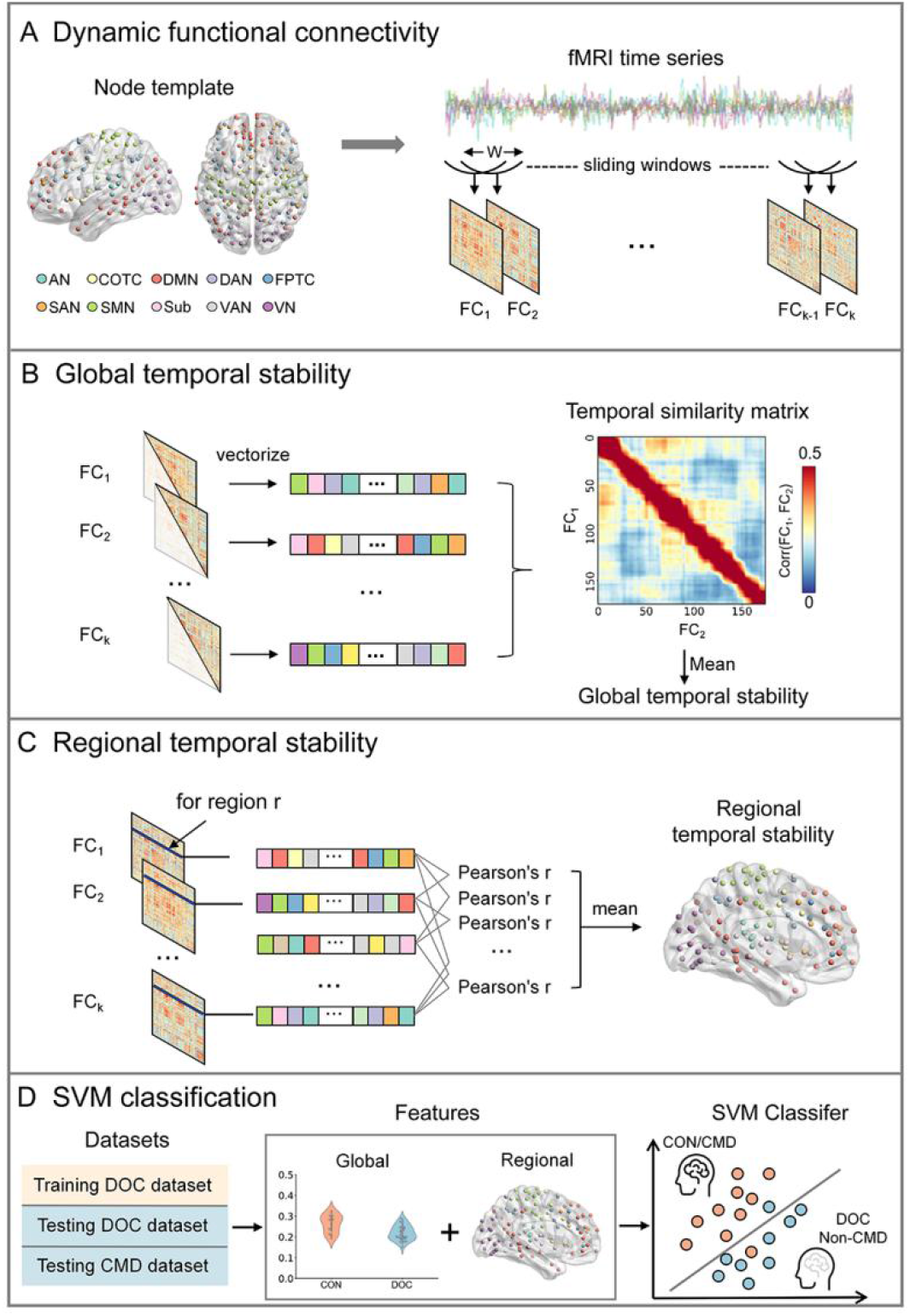
Pipeline of the Data Analysis. (A) To obtain the dynamic functional connectivity matrix, a sliding window approach (window length = 44 s, step = 2s) was used, where the Pearson’s correlation coefficient was calculated between each pair of brain regions based on a pre-defined brain template. Then the (B) global (i.e., whole-brain) and (C) regional temporal stability were defined as the similarity (using Pearson’s correlation) of the global and regional functional connectivity patterns across the time windows. (D) Finally, the temporal stability was used as features, and an SVM classifier was trained and evaluated in the first DOC dataset to investigate its performance in discriminating controls and DOC. The fully-trained SVM model was applied to the second independent dataset to test its generalizability. The model was further applied to the third independent dataset to test whether it could correctly identify patients with CMD. W = window length; FC = functional connectivity; AN = auditory network; COTC = cingulo-opercular task control network; DMN = default mode network; DAN = dorsal attention network; FPTC = fronto-parietal task control network; SAN = salience network; SMN = sensory/somatomotor network; Sub = subcortical network; VAN = ventral attention network; VN = visual network; CON = controls with full awareness; DOC = disorders of consciousness; CMD = cognitive motor dissociation; SVM = Support Vector Machine.

### Global And Regional Temporal Stability Analysis

Firstly, dynamic functional connectivity matrices were obtained using an overlapping sliding window method (window length = 22 TRs, step size = 1 TR, TR = 2 s). Specifically, for each subject, all BOLD time series were segmented into k overlapping time windows. Within each temporal window of 22 TRs, a pairwise 226 × 226 FC matrix was built by computing the Pearson correlation coefficient between the BOLD (Blood-oxygen-level-dependent) signals from a pair of ROIs. As a result, a set of FC matrices was obtained, i.e., [FC_1_, FC_2_, …, FC_k_]. 22 TRs (44s) of window length was chosen according to previous studies, which suggested that a window in the range of 30–60 s has sufficient time points to be robust enough in detecting the rs-fMRI FC fluctuation for a reliable network identification.^20^

Secondly, given a set of FC matrices [FC_1_, FC_2_, …, FC_k_] for each window, we introduced the measure of global temporal stability between each pair of FC matrices.^24,34^ Specifically, the Pearson correlation coefficient was used to measure spatial similarity between the upper-triangular parts of the FC matrices across time windows:

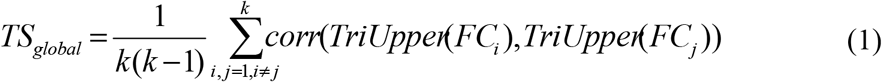

where TriUpper(FC_i_) stands for the upper triangle of the FC matrix at time window i, and corr(TriUpper(FC_i_), TriUpper(FC_j_)) stands for the correlation coefficient between two FC matrices. As illustrated in Fig. 1B, the temporal similarity matrix was firstly obtained, which depicted the spatial similarity between any pair of FCs, i.e., the stability of global FC pattern across time windows. The global temporal stability was then defined as the mean value of the off-diagonal elements of the temporal similarity matrix.

Finally, to characterize the temporal stability of the FC pattern associated with a given ROI, we introduced the measure of regional temporal stability that was slightly modified from recent studies.^25,35^ As shown in Figure 1C, the functional architecture of the region r across different time windows was extracted, which represented the overall FC pattern of region r across time windows, then the temporal stability of this region was defined as:

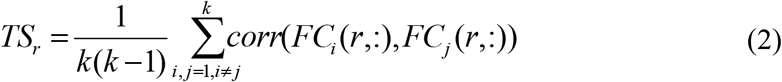

where FC_i_(r,:) stands for the FC pattern between the region r and all other regions at time window i, and corr(FC_i_(r,:), FC_j_(r,:)) stands for the correlation coefficient between two FC patterns. The regional temporal stability was then defined as the average correlation coefficient among all FC patterns of brain region r across time windows.

### Classification Analysis

Following the above procedures, a total of 227 features (1 global temporal stability and 226 regional temporal stability) were extracted for each participant. Prior to model training, all features were normalized into Z-scores. Then, as shown in Figure 1D, to test the predictive value of the calculated features (i.e., temporal stability) for classifying controls and DOC, a linear support vector machine (SVM) algorithm was applied using scikit-learn with default settings. Specifically, the SVM classifier were trained and evaluated on the dataset DOC-HS (n = 52) using a leave-one-subject-out (LOSO) cross-validation, where controls were labeled as 1, and DOC (including MCS/UWS) patients as 0. In each LOSO cross-validation fold, we used a method of SVM recursive feature elimination with cross-validation (SVM-RFECV, implemented in scikit-learn with default settings) to select the most informative features and reduce the risk of overfitting, in which model performance is iteratively tested with and without specific features.^36^ Furthermore, since the selected features are different in each LOSO cross-validation fold, we used the voting approach to choose features that occurred in at least 50% of the folds of cross-validation as the most important features.^35^ This yielded a smaller subset of 41 features of regional temporal stability (see Supplementary Figure 2). To test for robustness, using the selected features, we also evaluated whether the fully trained SVM model could generalize to the independent dataset DOC-ZJ (n = 56). Finally, to evaluate the capacity of the model to correctly identified DOC patients who was subsequently diagnosed as CMD, the same SVM model was also applied to the independent dataset CMD-GH/ZJ (n = 11). Specifically, a prediction was considered correct when a DOC patient was predicted as label 1 (as controls) and his/her subsequent BCI diagnosis was CMD, and vice versa.

### Statistical Analysis

For the difference of global and regional temporal stability, an independent two-sample t-test (two-sided) was applied between controls and DOC for dataset DOC-HS (16 CON vs. 36 DOC). To quantify the classification performance, receiver operating characteristic (ROC) analysis was used to calculate the area under the curve (AUC), which measures the ability of the SVM model in discriminating controls, CMD or DOC, and potential non-CMD. The value of the AUC ranges from 0 to 1, where 0 denotes 100% inaccurate predictions, an 1 denotes 100% accurate predictions. In addition, for each dataset, a confusion matrix was generated to measure results of the SVM classification. A χ^2^ test was then used to assess the statistical significance between the labels predicted by the classifier and the actual class labels. For the above statistic comparison, a Bonferroni correction at alpha < 0.05 was applied.

### Data Availability

The data that support the findings of the current study may be requested from the corresponding author upon reasonable request.

## Results

### Impaired Global Temporal Stability in Patients with DOC

Firstly, we investigated the difference of global temporal stability between controls and DOC, by measuring the similarity of their global FC pattern across different time windows. In Figure 2A, two examples were shown for visualization of the temporal similarity matrix (window length = 22 TRs, i.e., 44 s), one is a control (top row) and the other a DOC patient (bottom row). For each matrix, its elements represent the Pearson correlation coefficient of the global FC pattern between every pair of time windows. In general, temporal similarity of the DOC was markedly impaired, which reflected an unstable or unsynchronized global FC pattern across time windows.

**Figure 2.**
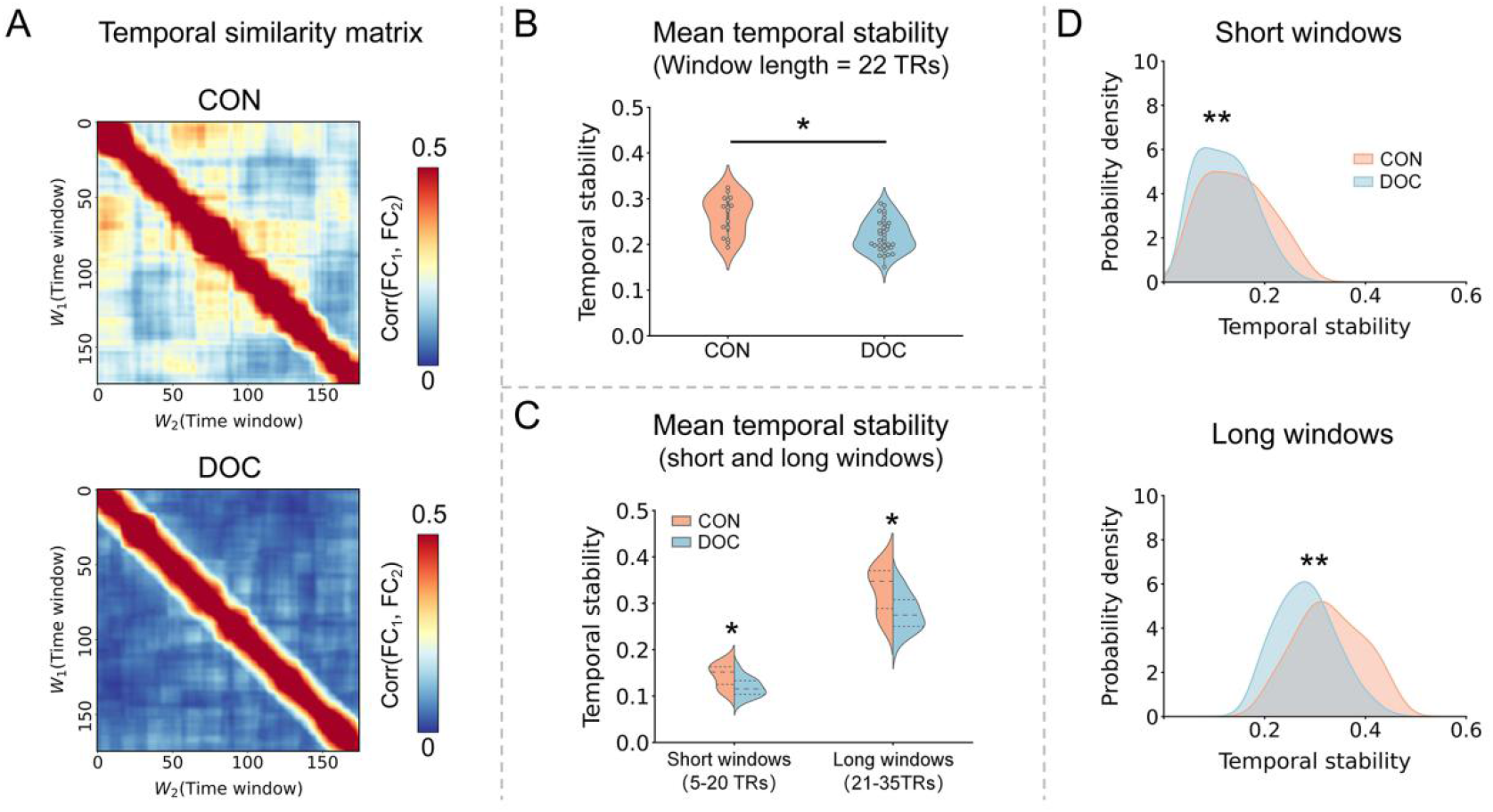
It should be noted that all results presented in the figure were obtained from the dataset DOC-HS. (A) Temporal similarity matrix for one representative control and DOC participant, respectively, which measures the similarity (using Pearson’s correlation) of the global (i.e., whole-brain) functional connectivity patterns across different time windows. (B) The group-level comparison of the global temporal stability between the DOC and control group was performed, where a window length of 22 TRs (i.e., 44s) was used. (C) The group-level difference was further validated for both short (mean value over window length from 5 to 20 TRs) and long window (mean value over window length from 21 to 35 TRs) ranges, (D) as well as their distribution. CON = controls with full awareness; DOC = disorders of consciousness; * means p < 0.05, ** means p < 0.01, Bonferroni corrected.

In addition, we quantified the difference of global temporal stability between the controls and DOC at a group-level, by averaging off-diagonal elements of each individual temporal similarity matrix. In the top row of Figure 2B, it can be seen that the global temporal stability was impaired in DOC relative to controls, at a window length of 22 TRs (two-sample t-test, p = 0.015). To validate the robustness of different window lengths, we performed the global temporal stability analysis at two different window ranges: a “short” range (from 5 to 20 TRs) and a “long” range (from 21 to 35 TRs). The results were consistent as the above, both in the mean value at the two window ranges (Figure 2C, two-sample t-test, p = 0.017 for “short” range, p = 0.015 for “long” range), and the distribution (Figure 2D, two-sample Kolmogorov-Smirnov test, p = 0.002 for “short” range, p < 0.001 for “long” range). All p values above were Bonferroni-corrected.

### Impaired Regional Temporal Stability in Patients with DOC

We further investigated the difference of regional temporal stability between controls and DOC, measuring the stability of FC pattern between the region and all other regions across time. Figure 3A shows the regional temporal stability (window length = 22 TRs) at a group level, where the stability value was significantly lower in DOC compared with controls, especially in the range of the DMN. This confirmed that the regional temporal stability in COTC, DMN, FPTC, SAN was significantly impaired in DOC patients (Figure 3B, p < 0.05, Bonferroni-corrected), and this finding is consistent at a network-level (Figure 3C, p value < 0.01, Bonferroni-corrected).

**Figure 3.**
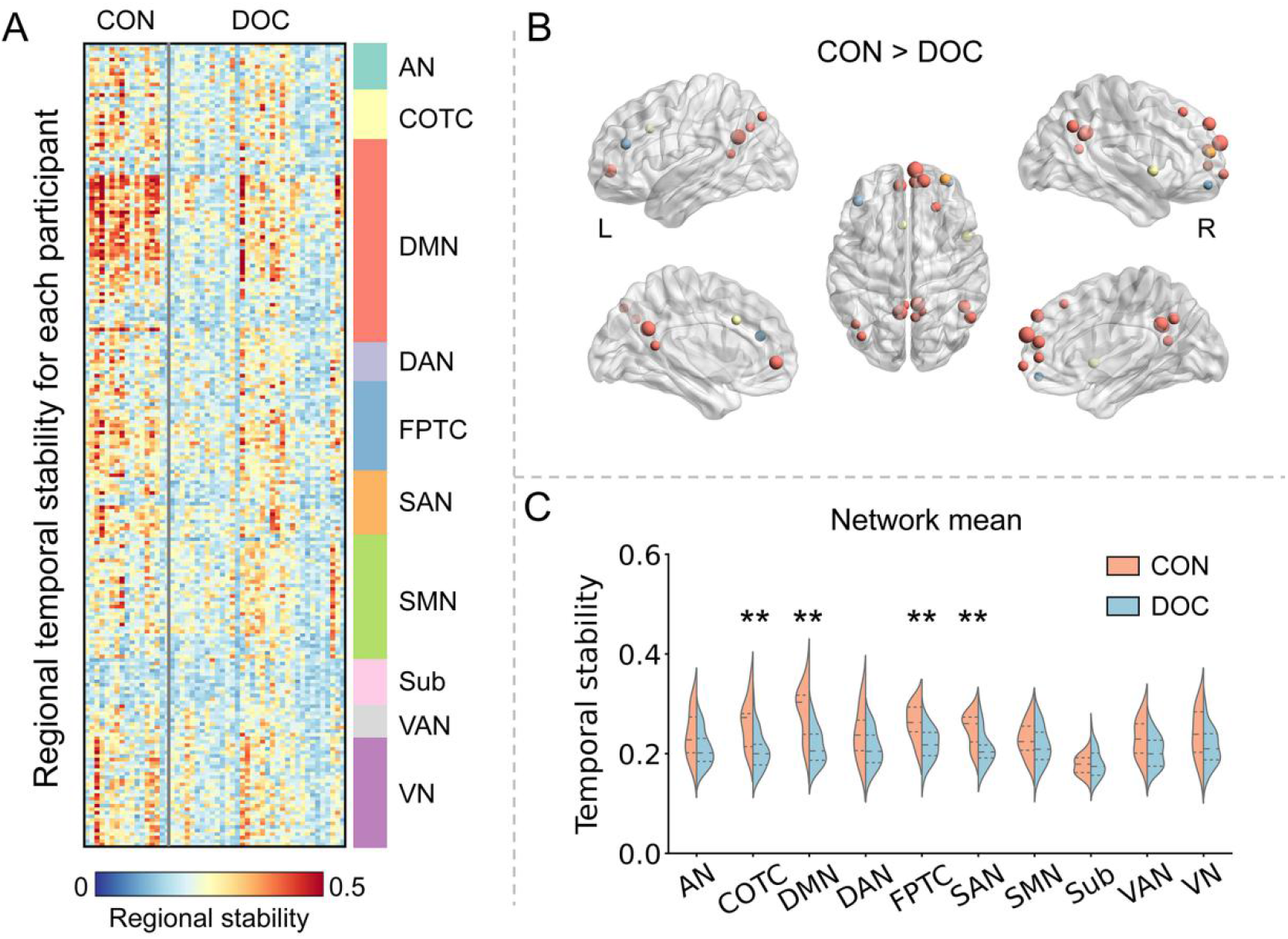
Comparison of Regional Temporal Stability Between Patients With DOC and Controls. It should be noted that all results presented in the figure were obtained from the dataset DOC-HS. (A) The regional temporal stability (window length = 22 TRs) value for each control and DOC participant across the ten functional networks. (B) Brain regions with significantly (p < 0.05 Bonferroni corrected) higher regional temporal stability for controls than the DOC patients, where the size and color of the node denotes the t-value and its corresponding network. (C) Group-level comparison of network mean temporal stability (by averaging the corresponding regions for each network). AN = auditory network; COTC = cingulo-opercular task control network; DMN = default mode network; DAN = dorsal attention network; FPTC = fronto-parietal task control network; SAN = salience network; SMN = sensory/somatomotor network; Sub = subcortical network; VAN = ventral attention network; VN = visual network; CON = controls with full awareness; DOC = disorders of consciousness; ** means p < 0.01, Bonferroni corrected.

### Correct Identification of CMD Using Temporal Stability

Finally, we trained an SVM classifier with selected features to test whether the model could discriminate controls and DOC in dataset DOC-HS, and validated it in the independent dataset DOC-ZJ. Additionally, the fully-trained model was further applied to the independent CMD-GH/ZJ dataset, to test its capacity in correctly identifying CMD from potential non-CMD. Specifically, the SVM recursive feature elimination with cross-validation (SVM-RFECV) method (implemented in scikit-learn with default settings) was used to select the most informative features, where 41 features of regional temporal stability were selected (see Supplementary Figure 2). In general, the SVM model showed a robust classification performance (accuracy = 90%) in discriminating controls and DOC in dataset in DOC-HS using a leave-one-subject-out cross-validation (Figure 4A), and showed good generalizability (accuracy = 82%) in the dataset DOC-ZJ (Figure 4B). More interestingly, as shown in Figure 4C, we also observed that the classifier generalized very well to the independent dataset CMD-GH/ZJ (Accuracy: 91%). Most importantly, all four DOC patients (3 UWS and 1 MCS at the time of rs-fMRI scanning) with a subsequent CMD diagnosis in BCI were correctly classified by the SVM model. Note that one DOC patient (being in UWS at the time of scanning) was misclassified as a control (i.e., being in fully conscious state), while his subsequent BCI diagnosis was potential non-CMD.

**Figure 4.**
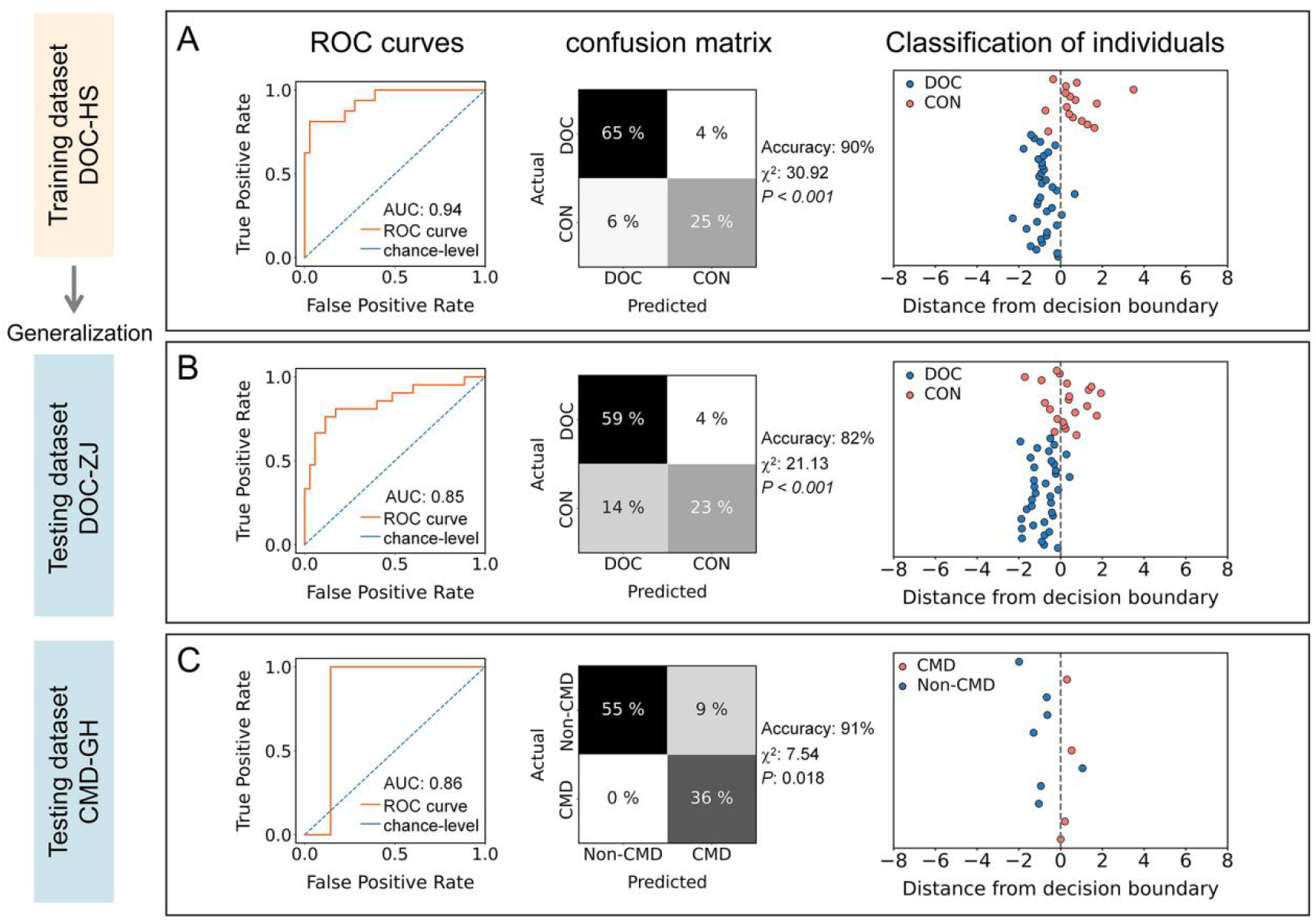
Classification Results of the Three Datasets. The temporal stability was used as features to build an SVM model, to test its performance in discriminating controls and DOC, as well as in identifying CMD. (A) The model was firstly trained and validated in the first dataset (DOC-HS) using a leaving-one-subject-out cross validation method, with which the ROC curve, confusion matrix, and individual classification outcome was displayed. (B) The fully-trained model was further applied to the independent dataset DOC-ZJ to test its generalizability. (C) The same model was then applied to the independent dataset CMD-GH/ZJ, to test its capacity in correctly identifying patients with CMD. ROC = receiver operating characteristic; AUC = area under curve; CON = controls with full awareness; DOC = disorders of consciousness; CMD = cognitive motor dissociation;

Specifically, in the left column of the Figure 4A-C, the area under the ROC curves (AUC) was calculated for dataset DOC-HS (AUC = 0.94), dataset DOC-ZJ (AUC = 0.85), and dataset CMD-GH/ZJ (AUC =0.86), respectively. The middle column of the Figure 4A-C plots the confusion matrix generated by the SVM for each dataset, in which a chi-squared test was used to estimate the classifier’s performance. The test showed consistently significant results for the dataset DOC-HS (χ^2^ = 30.92, p < 0.001), DOC-ZJ (χ^2^ = 21.13, p < 0.001), and CMD-GH/ZJ (χ^2^ = 7.54, p = 0.018), respectively. Finally, the right column of the Figure 4A-C showed the result of individual classification for each dataset, which depicted the proportional value of the distance for each individual from the separating hyperplane.

For validation purposes, we also performed the same classification analysis, using only the significant features (1/21 global/regional temporal stability) from dataset DOC-HS, and found a lowered classification performance (accuracy for dataset DOC-HS, DOC-ZJ, CMD-GH/ZJ: 73%, 70%, 82%). This suggests that choosing features according to its group-level significance might lose some meaningful information for classification, which is consistent with previous studies.^32,37^

## Discussion

The current study investigated the difference of temporal stability between fully conscious participants and patients with DOC, and tested its capacity in detecting covert awareness in CMD, whose awareness may be intact and yet unable to be detected with behavioral assessment. Our results showed impaired temporal stability in DOC, especially in the network of COTC, DMN, FPTC, and SAN. More importantly, using temporal stability as features, our classification analysis showed that the SVM model could not only discriminate DOC and controls with a high performance, but also able to correctly identify CMD. Overall, the current findings showed that consciousness was likely to rely on a stable functional architecture over time. Based on the current results, we believe that we could have developed an alternative way for diagnosing CMD patients that is highly convenient.

The most important finding of the current study is that the alteration of temporal stability could be informative in identifying CMD. Among the 11 DOC patients, our model correctly classified 10 patients, which was consistent with their subsequent BCI diagnosis, including 4 CMD and 6 potential non-CMD patients. In line with previous studies, our finding supported that a proportion of DOC patients may maintain covert awareness while fail to be detected with behavioral assessment.^9,13^ Current studies testing the patients’ awareness used either task-based fMRI (e.g., mental imagery tasks)^10^ or EEG-based BCI paradigms (e.g., item-selection task),^38^ both of which require several cognitive functions, e.g., working memory, and object recognition.^39^ As a result, some DOC patients diagnosed as potential non-CMD due to his/her deficit of memory or fluctuations of attention, could still maintain covert awareness. This issue was emphasized in a previous study that, a non-significant BCI accuracy was not definitive in proving absence of awareness (i.e., potential non-CMD), and false-negative findings in BCI experiments were possible,^11^ which could be the case for the one DOC patient (being in UWS at the time of fMRI scanning) misclassified by our model in the current study.

Furthermore, we found that the temporal stability in regions of several functional networks were impaired in DOC, including COTC, DMN, FPTC, and SAN, which highlights the importance of these networks in sustaining normal awareness. Among these functional networks, the DMN is involved in mind-wandering, and processing of the autobiographical and self-related information,^40^ and reduction in the functional connectivity in this network was consistently observed during neuropathological (e.g., DOC),^14,41^ pharmacological (e.g., propofol-induced sedation),^30,42^ and physiological (e.g., non-rapid eye movement sleep) unconscious states.^43^ The FPN and COTC on the other hand, are two major task-positive functional networks, which were found associated with goal-directed behaviors and perceptual processing,^44^ thus playing a central hub-like role in supporting human consciousness.^45,46^ The SAN, which has been implicated in interoceptive information processing^47^, has also been found to be reduced in its functional connectivity in multiple unconscious states.^48,49^ Beyond the existing evidence, our results showed that DOC suffered from a loss of stable functional architecture across time, reflected by an impaired temporal stability, which could mean that a fully conscious states might be characterized by the opposite, i.e., maintaining a stable functional organization over time. This assumption was supported by a previous study, where functional connectivity during DOC and anesthesia states exhibited a significant reduction in the temporal states of high integration,^29^ suggesting that unconscious states could be dominated by a fragmentized and unstable (i.e., with a low temporal integration) functional architecture. In summary, our findings highlighted the importance of temporal stability in supporting consciousness.

Several issues should be noted. Firstly, the sample size of DOC patients who underwent the rs-fMRI and BCI experiment was relatively small (n = 11). Although our classification results exhibited a good performance (10 of 11 patients were correctly classified), showing a potential of rs-fMRI in detecting covert awareness in the DOC, more studies in the future are needed to further test its robustness. Secondly, in the current study, the rs-fMRI scanning and BCI assessment were not performed at the same time (mean time interval between rs-fMRI and BCI assessment: 116 days), and it was possible that patients’ conscious states may vary between the two assessments. Although, among the 11 patients, 9 patients had consistent behavioral diagnosis (i.e., Coma Recovery Scale-Revised, CRS-R scale) between the time of rs-fMRI and the BCI assessment, as well as consistent SVM classification outcomes with the BCI diagnosis. Furthermore, even the two patients who were behaviorally diagnosed as UWS (at the time of rs-fMRI scanning) and emerged to MCS (at the time BCI assessment), were both classified as having awareness using the resting-state temporal stability. It is therefore possible, that these two patients might had covert awareness, which failed to be detected by the behavioral assessment.^7,9,18^ Thirdly, although participants were asked to keep their eyes open before the experiment, we cannot fully rule out the possibility that some of the participants may felt drowsy and slept during the scanning,^50^ which might lead to some fully conscious participants (i.e., controls) misclassified as DOC in the current study.

In conclusion, our results exhibited that the temporal stability was impaired in patients with DOC. Specifically, patients with DOC were accompanied by a reduction of temporal stability in several functional networks, including COTC, DMN, FPTC, and SAN. More interestingly, we built an SVM model that used temporal stability as features, and showed a good performance in correctly identifying potential CMD. Taken together, the current study offered new insight into the neural mechanism of consciousness, and showed a potential diagnostic value of rs-fMRI for CMD patients.

## Abbreviations

CRS-R: Coma Recovery Scale-Revised
DOC: disorders of consciousness
SVM: Support Vector Machine
UWS: unresponsive wakefulness syndrome
MCS: minimally conscious state
CMD: cognitive motor dissociation
dFC: dynamic functional connectivity
rs-fMRI: resting-state fMRI
BCI: brain-computer interface.

## Acknowledgments

We thank all the patients and healthy participants for taking part in this study. We also thank Mingxia Wang for her help in improving the English in the manuscript.

## Funding

This work was supported by Key Realm R&D Program of Guangzhou (202007030005), the National Natural Science Foundation of China (31971032, 8197414, 82171174), the Major Program of the National Social Science Fund of China (18ZDA293), the Basic and Applied Basic Research Foundation of Guangdong Province (2020A1515011250), Guangdong-Hong Kong-Macao Greater Bay Area Center for Brain Science and BrainInspired Intelligence Fund (2019023), Key Realm R&D Program of Guangdong Province (2019B030335001).

## Author contributions

**Hang Wu**: Methodology, Software, Formal Analysis, Writing - Original Draft, Writing - Review & Editing, Visualization. **Qiuyou Xie**: Investigation, Resources, Writing - Review & Editing; **Jiahui Pan**: Investigation, Writing - Review & Editing; **Qimei Liang**: Resources; **Yue Lan**: Resources; **Yequn Guo**: Investigation; **Junrong Han**: Writing - Review & Editing; **Musi Xie**: Visualization; **Yueyao Liu**: Writing - Review & Editing; **Liubei Jiang**: Software; **Xuehai Wu**: Investigation, Writing - Review & Editing; **Yuanqing Li**: Methodology, Writing - Review & Editing; **Pengmin Qin**: Conceptualization, Methodology, Writing - Review & Editing, Supervision.

## Competing interests

The authors report no competing interests.

## Supplementary material

**Supplementary Table 1.** Detailed Demographic and Clinical Information for Dataset DOC-HS

| Patient<br>Number | Age | Sex | Aetiology | Days Since<br>Injury | CRS-R | Diagnosis |
| --- | --- | --- | --- | --- | --- | --- |
| P1 | 52 | M | TBI | 210 | 6 | UWS |
| P2 | 47 | M | SIH | 315 | 6 | UWS |
| P3 | 35 | M | SIH | 300 | 4 | UWS |
| P4 | 46 | M | TBI | 168 | 1 | UWS |
| P5 | 53 | M | TBI | 30 | 3 | UWS |
| P6 | 26 | M | TBI | 143 | 6 | UWS |
| P7 | 45 | M | TBI | 12 | 6 | UWS |
| P8 | 67 | F | TBI | 10 | 3 | UWS |
| P9 | 38 | M | TBI | 81 | 3 | UWS |
| P10 | 35 | F | TBI | 21 | 5 | UWS |
| P11 | 17 | M | TBI | 16 | 6 | UWS |
| P12 | 46 | M | TBI | 25 | 2 | UWS |
| P13 | 62 | M | TBI | 24 | 4 | UWS |
| P14 | 57 | F | TBI | 15 | 3 | UWS |
| P15 | 57 | M | SIH | 33 | 2 | UWS |
| P16 | 39 | F | TBI | 31 | 6 | UWS |
| P17 | 46 | M | SIH | 18 | 2 | UWS |
| P18 | 46 | M | TBI | 162 | 5 | UWS |
| P19 | 44 | M | TBI | 21 | 3 | UWS |
| P20 | 52 | F | SIH | 51 | 4 | UWS |
| P21 | 43 | F | SIH | 162 | 8 | UWS |
| P22 | 39 | M | SIH | 221 | 4 | UWS |
| P23 | 46 | M | Anoxic | 23 | 3 | UWS |
| P24 | 51 | M | TBI | 100 | 4 | UWS |
| P25 | 60 | M | SIH | 109 | 6 | UWS |
| P26 | 24 | M | Anoxic | 5 | 7 | UWS |
| P27 | 16 | M | TBI | 4 | 12 | MCS |
| P28 | 56 | M | TBI | 18 | 8 | MCS |
| P29 | 48 | M | TBI | 19 | 10 | MCS |
| P30 | 18 | M | TBI | 30 | 6 | MCS |
| P31 | 32 | F | TBI | 73 | 12 | MCS |
| P32 | 17 | M | TBI | 44 | 12 | MCS |
| P33 | 36 | M | TBI | 82 | 13 | MCS |
| P34 | 44 | M | TBI | 98 | 8 | MCS |
| P35 | 64 | M | SIH | 157 | 8 | MCS |
| P36 | 30 | M | TBI | 240 | 8 | MCS |
| P37 | 54 | F | TBI | 4 | 23 | BI |
| P38 | 34 | M | TBI | 4 | 23 | BI |
| P39 | 24 | F | TBI | 139 | 23 | BI |
| P40 | 52 | F | TBI | 137 | 23 | BI |
| P41 | 29 | F | TBI | 6 | 23 | BI |
| P42 | 18 | M | TBI | 182 | 23 | BI |
| P43 | 46 | M | TBI | 43 | 23 | BI |
| P44 | 50 | F | SIH | 9 | 23 | BI |
| P45 | 49 | M | TBI | 57 | 23 | BI |
| P46 | 18 | M | TBI | 105 | 23 | BI |
| P47 | 33 | F | SIH | 4 | 23 | BI |
| P48 | 70 | M | TBI | 11 | 23 | BI |
| P49 | 46 | M | SIH | 41 | 23 | BI |
| P50 | 64 | M | SIH | 30 | 23 | BI |
| P51 | 21 | M | TBI | 621 | 23 | BI |
| P52 | 24 | M | TBI | 165 | 23 | BI |

Note: UWS = unresponsive wakefulness syndrome; MCS = minimally conscious state; BI = fully conscious participants with brain injury history; CRS-R = Coma Recovery Scale-Revised; SIH = spontaneous intracerebral hemorrhage; TBI = traumatic brain injury.

**Supplementary Table 2.** Detailed Demographic and Clinical Information for Dataset DOC-ZJ

| Patient<br>Number | Age | Sex | Aetiology | Days Since<br>Injury | CRS-R | Diagnosis |
| --- | --- | --- | --- | --- | --- | --- |
| P1 | 60 | F | TBI | 25 | 4 | UWS |
| P2 | 53 | M | SIH | 179 | 2 | UWS |
| P3 | 56 | M | Anoxic | 55 | 6 | UWS |
| P4 | 42 | M | TBI | 42 | 1 | UWS |
| P5 | 27 | F | Anoxic | 25 | 7 | UWS |
| P6 | 32 | F | Anoxic | 75 | 5 | UWS |
| P7 | 56 | F | TBI | 73 | 4 | UWS |
| P8 | 39 | M | Anoxic | 48 | 2 | UWS |
| P9 | 18 | M | Anoxic | 56 | 6 | UWS |
| P10 | 39 | M | SIH | 97 | 1 | UWS |
| P11 | 48 | M | SIH | 110 | 4 | UWS |
| P12 | 36 | F | Anoxic | 62 | 6 | UWS |
| P13 | 13 | M | TBI | 35 | 6 | UWS |
| P14 | 50 | F | Anoxic | 32 | 5 | UWS |
| P15 | 34 | M | TBI | 38 | 6 | UWS |
| P16 | 67 | F | Anoxic | 45 | 5 | UWS |
| P17 | 35 | F | Anoxic | 62 | 6 | UWS |
| P18 | 40 | M | Anoxic | 67 | 8 | UWS |
| P19 | 39 | M | SIH | 116 | 3 | UWS |
| P20 | 22 | M | TBI | 47 | 5 | UWS |
| P21 | 46 | M | SIH | 62 | 3 | UWS |
| P22 | 58 | M | TBI | 153 | 8 | UWS |
| P23 | 55 | M | TBI | 195 | 6 | UWS |
| P24 | 69 | F | Anoxic | 35 | 5 | UWS |
| P25 | 64 | M | TBI | 71 | 3 | UWS |
| P26 | 55 | F | Anoxic | 37 | 3 | UWS |
| P27 | 52 | M | Anoxic | 34 | 10 | MCS |
| P28 | 30 | F | TBI | 78 | 7 | MCS |
| P29 | 23 | M | SIH | 31 | 8 | MCS |
| P30 | 36 | M | TBI | 89 | 15 | MCS |
| P31 | 46 | F | SIH | 115 | 12 | MCS |
| P32 | 58 | M | SIH | 29 | 11 | MCS |
| P33 | 49 | F | SIH | 42 | 7 | MCS |
| P34 | 20 | M | TBI | 287 | 10 | MCS |
| P35 | 53 | F | TBI | 31 | 7 | MCS |

Note: UWS = unresponsive wakefulness syndrome; MCS = minimally conscious state; CRS-R = Coma
Recovery Scale-Revised; SIH = spontaneous intracerebral hemorrhage; TBI = traumatic brain injury;
“-” means not applicable.

**Supplementary Table 3.** Detailed Demographic and Clinical Information for Dataset CMD-GH/ZJ

| Patient<br>Number | Age | Sex | Aetiology | Days Since<br>Injury | CRS-R at<br>rs-fMRI | Diagnosis at<br>rs-fMRI | CRS-R<br>at BCI | Diagnosis<br>at BCI | BCI<br>Accuracy |
| --- | --- | --- | --- | --- | --- | --- | --- | --- | --- |
| P1 | 48 | M | Anoxic | 19 | 3 | UWS | 7 | UWS | **66** |
| P2 | 39 | M | Anoxic | 67 | 5 | UWS | 5 | UWS | **72** |
| P3 | 39 | M | TBI | 7 | 5 | UWS | 6 | UWS | 52 |
| P4 | 46 | M | TBI | 73 | 5 | UWS | 5 | UWS | 62 |
| P5 | 51 | M | TBI | 45 | 6 | UWS | 11 | MCS | 52 |
| P6 | 50 | M | SIH | 106 | 3 | UWS | 9 | MCS | **66** |
| P7 | 58 | M | Anoxic | 140 | 5 | UWS | 5 | UWS | 52 |
| P8 | 15 | F | Anoxic | 33 | 12 | MCS | 12 | MCS | 60 |
| P9 | 35 | M | TBI | 42 | 6 | UWS | 6 | UWS | 62 |
| P10 | 56 | F | Anoxic | 208 | 12 | MCS | 12 | MCS | **66** |
| P11 | 58 | M | TBI | 117 | 8 | MCS | 9 | MCS | 58 |

Note: UWS = unresponsive wakefulness syndrome; MCS = minimally conscious state; CMD = cognitive motor dissociation; BCI = brain-computer interface; CRS-R = Coma Recovery Scale-Revised; SIH = spontaneous intracerebral hemorrhage; TBI = traumatic brain injury; “-” means not applicable; BCI accuracies significantly higher than chance level (i.e., potential DOC patients with CMD) are highlighted in bold.

**Supplementary Figure 1.**
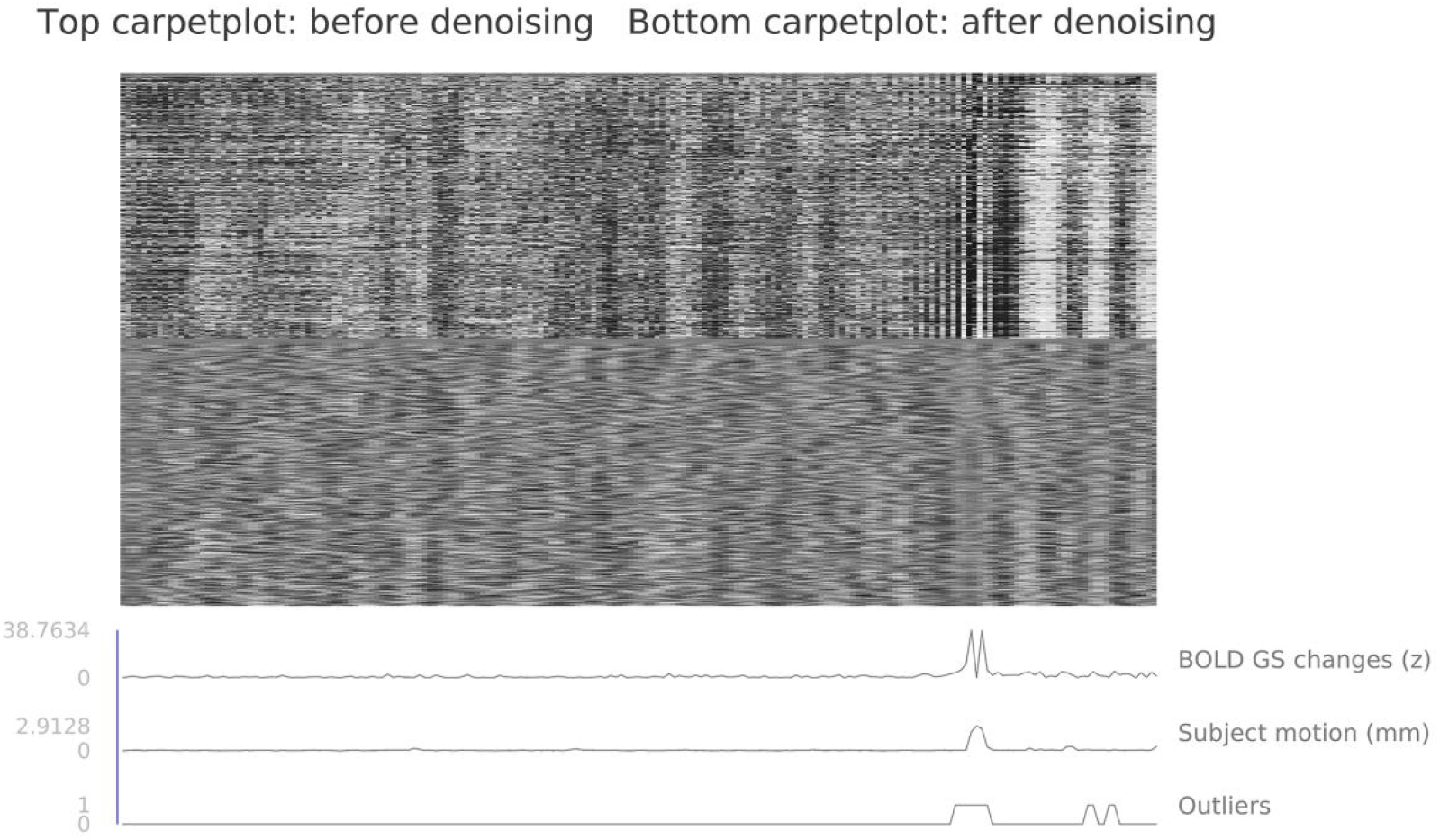
Illustration for the Effect of Denoise Procedure on an Example DOC Participant. The two carpetplots show the voxel-wise time series before (top one) and after (bottom one) the denoise procedure, where the corresponding bottom lines show the outlier time points (i.e., extreme global signal and motion value) detected by the ART toolbox.

**Supplementary Figure 2.**
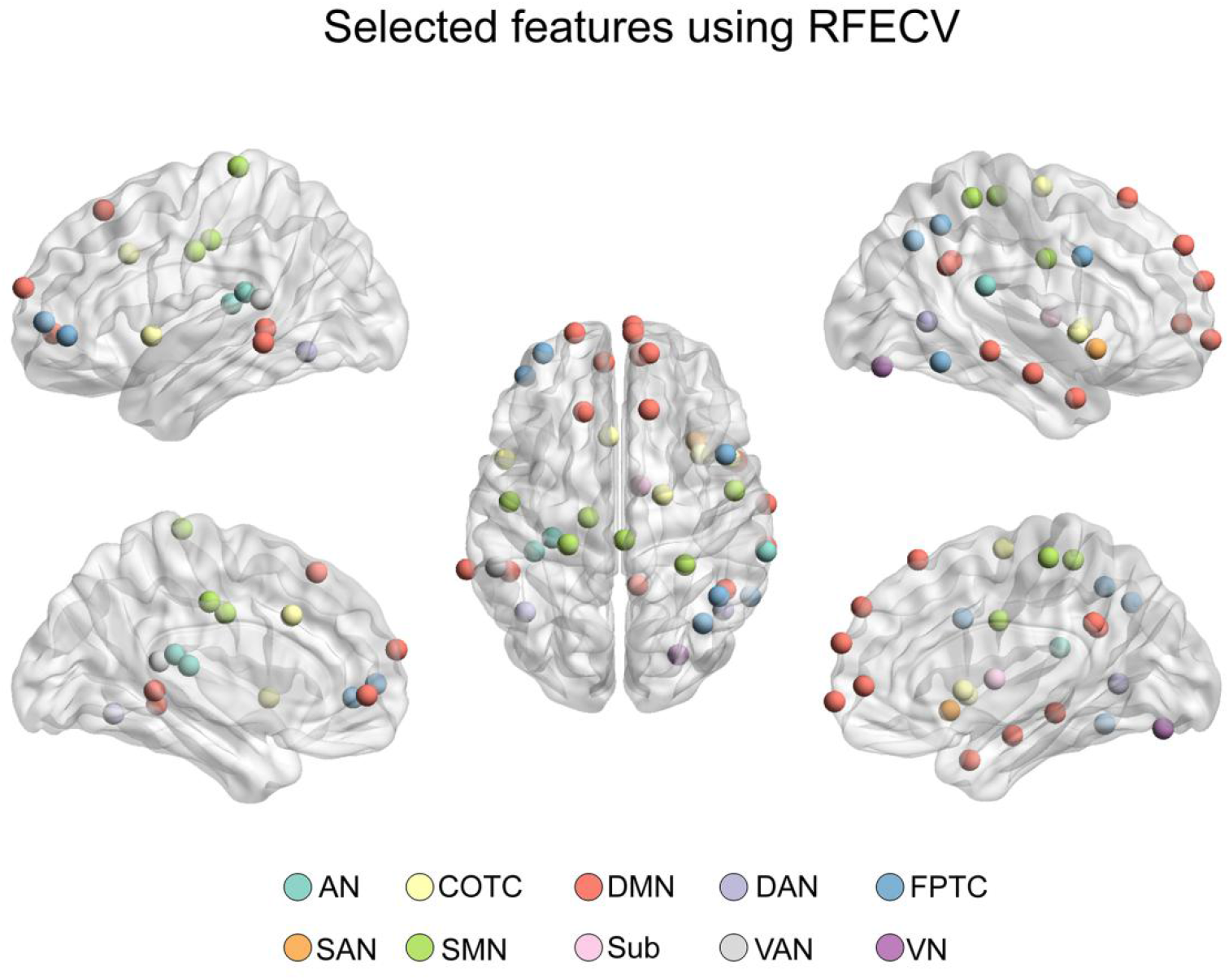
The most Important Features After Feature Selection. The brain nodes denote the regional temporal stability selected as important features with the SVM recursive feature elimination with cross-validation (SVM-RFECV) method. The color of each node denotes their corresponding functional networks. AN = auditory network; COTC = cingulo-opercular task control network; DMN = default mode network; DAN = dorsal attention network; FPTC = fronto-parietal task control network; SAN = salience network; SMN = sensory/somatomotor network; Sub = subcortical network; VAN = ventral attention network; VN = visual network.

## Notes

### Competing Interest Statement

The authors have declared no competing interest.

